# Heterogeneity of synaptic connectivity in the fly visual system

**DOI:** 10.1101/2023.08.29.555204

**Authors:** Jacqueline Cornean, Sebastian Molina-Obando, Burak Gür, Annika Bast, Giordano Ramos-Traslosheros, Jonas Chojetzki, Lena Lörsch, Maria Ioannidou, Rachita Taneja, Christopher Schnaitmann, Marion Silies

## Abstract

Visual systems are homogeneous structures, where repeating columnar units are stereotypically arranged to retinotopically cover the visual field. Each of these columns contain many of the same neuron types that are distinguished by anatomic, genetic and – generally – by functional properties. However, there are exceptions to this rule. In the 800 visual columns of the *Drosophila* eye, there is an anatomically and genetically identifiable cell type with variable functional properties, Tm9. Since anatomical connectivity shapes functional neuronal properties, we identified the presynaptic inputs of several hundred Tm9s across both optic lobes using the FAFB connectome dataset and FlyWire analysis. Our work shows that Tm9 has three major, stereotypic, and many weaker, sparsely distributed inputs. This differs from the presynaptic connectivity of neurons with uniform properties, Tm1 and Tm2, which have only one major, and more stereotypic inputs than Tm9. Within the heterogeneous circuit architecture, we identified specific motifs, such as a set of wide-field neurons, which can be the source of the variable Tm9 physiology. Genetic synapse labeling combined with expansion microscopy showed that the heterogeneous wiring exists across individuals. Together, our data argue that the visual system uses heterogeneous, distributed circuit properties to achieve robust visual processing.

## Introduction

Neuronal connectivity in the brain can be remarkably precise. This is especially true for visual systems, which follow a similar blueprint of being highly homogeneous and structured networks. Here, neuronal cell types are stereotypically organized in parallel visual columns, which retinotopically cover visual space. This repetitive, mosaic-like structure ensures the parallel processing of visual features across the visual system. Each part of the visual system can extract the same information across visual space, including information about luminance or color contrast, edges, or the direction of local motion ^1–4^. Along the proximodistal axis of the visual system, visual neuron projections are organized into layers according to their cell type and function, with ON and OFF pathway neurons projecting to different layers in both the vertebrate and invertebrate visual systems ^4–7^. Within these structures, the homogeneous connectivity of individual cell types is considered a common design principle of diverse visual systems. As a consequence, cells of one type are considered anatomically, genetically, and functionally uniform ^6,8–10^.

Although this highly regular organization generalizes across visual animals, some aspects of the eye structure deviate from this homogeneity. For example, a specialized area in the eye of many insects detects skylight polarization, the dorsal rim area ^11,12^. In this area, the opsin composition in photoreceptors is different than in the rest of the eye ^11,13^. Some insects have areas in their eyes that are specialized for detecting mates or prey with enlarged ommatidia and mostly different downstream circuits^14^. Regional specializations are also found in the vertebrate retina. For example, the upper visual fields of mice and zebrafish larvae are strongly UV-sensitive ^15–17^, matching the statistics of their environment. Similarly, acute zones, or regions with higher retina ganglion cell density, vary in shape and size between species or even animals of the same species but living in different habitats, adapted to very finely sample important parts of their visual scene (reviewed in ^17^). All of these specializations across the eye have in common that they are “orderly” and tuned to the specific visuo-ecological niche of the animal. However, some stochasticity also exists in the patterning of the eye. In particular, different classes of photoreceptors involved in processing color or UV stimuli are stochastically distributed in humans and many other animals, forming retinal mosaics (reviewed in ^11,18^). Careful developmental studies and recent connectomics analysis in *Drosophila* have shown that this pattern is followed by ommatidia-type specific downstream circuitry, although the downstream neurons are then stereotypic for each specific ommatidial subtype ^19–21^. The visual pathways that process non-color stimuli and take input from photoreceptors expressing a single rhodopsin Rh1 are considered highly homogeneous throughout the visual system ^22^. No heterogeneity of circuit wiring across the fly eye has been described for these pathways.

However, some evidence argues for heterogeneity of synaptic wiring in the fly visual system. First, developmental processes set the stage for stochastic brain wiring in the fly eye: Already in photoreceptors, the growth cones of developing photoreceptors show stochastic filopodial dynamics, which in turn can affect wiring specificity ^23,24^. These filopodial dynamics depend on cell-autonomous processes and also on external developmental temperature ^25,26^, arguing that brain wiring might differ between individual units of the same eye, as well as between individual animals growing up under different environmental conditions. Second, deeper within visual pathways, a class of visual projection neurons shows wiring variability between individuals, which manifests in inter-individual behavioral variability ^27,28^. Third, variable physiological properties have been described in Tm9, a second order visual interneuron ^29^. Tm9 neurons are part of the OFF pathway, providing critical input to the direction-selective T5 neurons (Fig. 1a) ^29–33^. Tm9 is a cell type by most criteria: it is a columnar neuron with easily distinguishable morphology, it tiles the visual system, it expresses specific genetic markers (Fig. 1b), and is retinotopically organized ^5,31,33^. At the same time, Tm9 neurons show variability in their spatial receptive field sizes ^29^, as well as in their temporal response properties to visual stimuli (Fig. 1c). Where these variable functional properties of a peripheral visual system cell type originate from is not known.

**Fig. 1:**
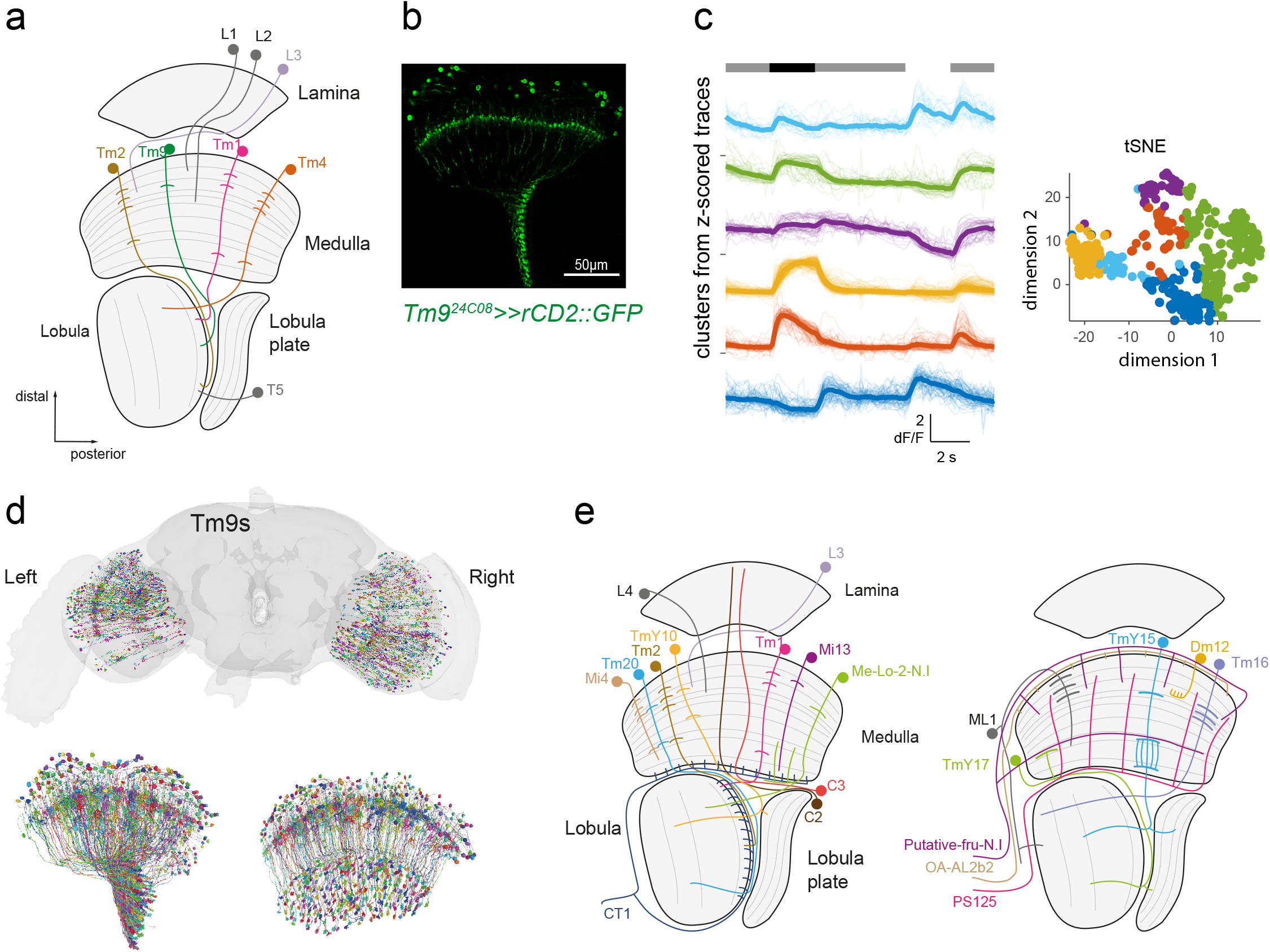
Mapping all inputs to Tm9 in FlyWire. **a)** Schematic of the fly visual system, including core circuitry of the OFF pathway. b) Confocal image of a visual system in which a Tm9-specific Gal4 line drives expression of GFP (green). Scale bar is 50 μm. c) In vivo calcium signals in Tm9 in response to ON-OFF-full field flashes. Shown are time traces (all individual traces and the mean) and tSNE analysis of Z-scored traces, color-coded upon k-means clustering (6 clusters). d) Overview of all 320 Tm9 neurons of the FAFB dataset analyzed in FlyWire (top), two views of all Tm9s analyzed in the right optic lobe. e) Schematic displaying the 20 input partners identified in this analysis to be present in at least 5 % of all columns.

Physiological properties are shaped by intrinsic properties as well as synaptic properties. Motivated by the variable responses to visual stimuli observed in Tm9, we explored the hypothesis that there is heterogeneity in Tm9’s presynaptic connectivity. Detailed synaptic connectivity based on EM analyses of the *Drosophila* optic lobes has been previously described, and contributed to our understanding how visual computations are implemented in peripheral visual circuitry ^3,22,22,32,34–37^. However, these efforts were restricted to individual or few columns. A recent, large dataset that covered most regions of the central brain - hemibrain ^38^ - excluded most of the visual system. The full adult female brain (FAFB) ^39^ is the first EM dataset that contains the two optic lobes, and together with state-of-the art connectome analysis (companion papers: ^40,41^) allows answering biological questions that require an analysis of connectivity across the entire visual system.

We here investigated the wiring variability of Tm9 presynaptic inputs in the FAFB connectome and using FlyWire analysis ^40,42^. We reconstructed several hundred Tm9 neurons and their presynaptic partners across both optic lobes. This work showed that three of the ∼ 20 presynaptic inputs are connected to Tm9 in each column, whereas the large majority of presynaptic inputs are sparsely connected to Tm9. Comparison of the presynaptic circuit architecture of Tm9 and two other Tm neuronal cell types shows that this degree of heterogeneity is Tm9 specific. Our analysis revealed circuit motifs that comprise different neurons, whose relative contribution, or variable connectivity can account for functional differences in Tm9 properties. Despite these functional motifs, there was no apparent organizational principle of Tm9 connectivity patterns across the eye, or between eyes, arguing that the spatial distribution of connectivity of many presynaptic inputs to Tm9 is stochastic. This example of heterogeneous wiring across the visual system suggests a structural basis for functional variation in visual processing across the eye, suggesting that visual system function tolerates or even benefits from a distribution of labor across the eye.

## Results

### Systematic reconstruction of presynaptic inputs to Tm9 across the Drosophila optic lobes

The FAFB dataset allows, for the first time, to comprehensively study wiring variation across the many parallel units of the visual system. To understand if variable physiological properties in a visual cell type can be accounted for by heterogeneity in its presynaptic inputs, we identified 320 Tm9s in the two optic lobes (n = 150 Tm9s left, 170 Tm9s right) (Fig. 1d). To analyze the presynaptic circuitry of Tm9 neurons across the eye, we mapped Tm9 postsynapses and their corresponding presynaptic sites using the Buhmann algorithm ^43^. We thus identified 35,100 synapses and subsequently reconstructed all fragments or neurons with at least 3 synapses to Tm9 (with help from many FlyWirers, see Supplementary Table 1). This corresponds to 2,724 cells, which accounted for 74.2 ± 9.5 % of all inputs to Tm9 across the two hemispheres. More detailed, manual proofreading identified further twigs of Tm9 dendrites, but did not significantly alter the identification of inputs as compared to backbone proofreading only (Extended Data Fig. 1a), arguing that our analysis reveals the full input architecture of Tm9. We then annotated the identity of 2,635 presynaptic cells (96.7 %) based on visual comparison with previous work ^5,19,36,37,44–46^, or by comparison with hemibrain ^38,47^. The repetitive organization of the visual system further allowed to assign neurons or groups of neurons into classes (cell types) based on similarity in the layer-specific arborization of cells. This way, almost all neurons that projected within the visual system could unambiguously be matched to previously described cell types (Extended Data Fig. 1b). Previously undescribed neurons were grouped based on their projection patterns. Across the 320 columns that we analyzed across the two visual systems, 20 different cell types gave presynaptic input to Tm9 in at least 5 % of the columns (Fig. 1e, Extended Data Fig. 1b). This included columnar and multi columnar neurons of the visual system, but also large neurons covering the entire optic lobe and projecting to or receiving projections from the central brain.

### Tm9 has many distributed presynaptic inputs

A first overview of the dataset showed that many presynaptic cells were variable and did not give input to Tm9 in all columns. Although 20 different cell types were presynaptic to Tm9, only some of these cell types were present as a presynaptic input of an individual Tm9 neuron (Fig. 2a). Specifically, Tm9 made a total of 78.1 ± 24.6 postsynaptic contacts (73.5 ± 25.4 synapses right and 83.3 ± 22.6 synapses left). These contacts belonged to an average of 8.2 ± 2.8 presynaptic cells per Tm9 neuron (8.2 ± 2.8 cells right and 8.3 ± 2.7 cells left). Because in some columns, several cells of one type connected on Tm9, this in turn accounted for 7.5 ± 2.3 presynaptic cell types per Tm9 neurons (7.4 ± 2.4 cells right and 7.5 ± 2.1 cells left) (Extended Data Fig. 2a). Remarkably, only three cell types were non-variable inputs to Tm9 (Fig. 2a). The cells that were presynaptic to Tm9 in all columns included two cells which gave presynaptic input to Tm9 in its dendritic region in the distal medulla: the well known presynaptic Tm9 lamina neuron input L3 ^31,48^, and the medulla intrinsic neuron Mi4, which has been extensively described as a component of the core circuitry for motion detection in the ON pathway ^37,49^(Fig. 2b,c). Furthermore, one of the homogeneous inputs to Tm9, CT1, synapses onto Tm9 axon terminals in each column. CT1 is a large, wide-field neuron with projections into both the lobula and medulla that forms highly compartmentalized projections in each of the visual columns, such that each projection can be treated as a columnar neuron ^32,37,50,51^ (Fig. 2b,c). We consider these as the core circuit motif of Tm9 connectivity (Fig. 2b).

**Fig. 2:**
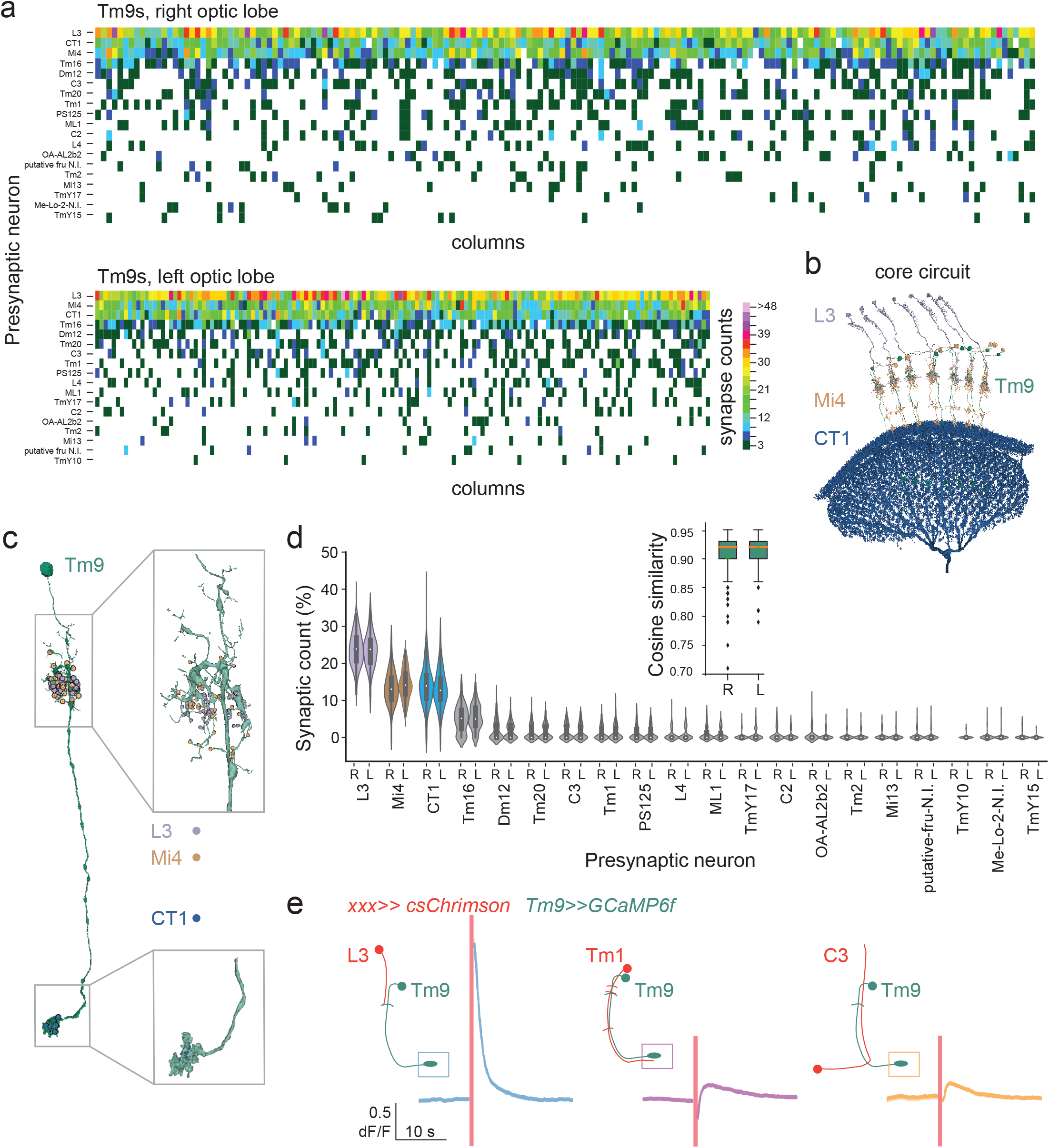
Analysis of heterogenous Tm9 inputs. **a)** Matrices showing synaptic counts of presynaptic inputs to Tm9 neurons of 170 columns and 150 columns of the right and left optic lobes, as long as they are present in >5 % of columns. **b)** One Tm9 (green) and its common presynaptic inputs L3 (purple), Mi4 (brown), and CT1 (blue). **c)** Spatial locations of the presynapses of L3 and Mi4 on the Tm9 dendrites, and CT1 synapses on the Tm9 axon terminal. **d)** Violin plots of the relative synaptic counts of Tm9 inputs, showing the median (white dot), the interquartile range (thick gray bar), 1.5x inter-quartile range (thin gray line) and outliers. Neurons with > 10 % contribution to synapse count are color-coded. Inset: Cosine similarity compared connectivity within the right (R) and left (L) optic lobes. **e)** In vivo calcium signals recorded in Tm9 neurons expressing GCaMP6f upon optogenetic activation of csChrimson expressed in L3 (n = 569 ROIs, 8 flies), Tm1 (n = 154 ROIs, N = 2 flies), or C3 (n = 658 ROIs, 8 flies), imaged in a *norpA* mutant background. The red line denotes

The three presynaptic inputs to Tm9 that were present in all columns also contributed with the highest count of synapses (Fig. 2d). L3 contributed the most with 25 ± 7.3 synapses per column, followed by Mi4 with on average 13.3 ± 7.0 and CT1 with 15.0 ± 5.7 synapses. However, these three cell types together still only contributed half (51.4 ± 6.4 % total, 51.4 ± 6.8 % right and 51 ± 6.3 % inputs left) of all presynaptic inputs to Tm9 (Fig. 2d). This suggests that the remaining, variable inputs which are not present in all columns, are not just the noise floor of all synaptic inputs. Each of the variable synaptic inputs to Tm9 only contributed roughly 5-10 % of all synaptic counts to one Tm9, arguing that all remaining, variable inputs are highly distributed (Fig. 2d). The synaptic input architecture between Tm9s from the right or the left optic lobe did not vary (Fig. 2d). We next tested functionally if the variable inputs, which have lower synapse counts, can significantly contribute to the physiological properties of Tm9. We optogenetically activated the major Tm9 input L3 as a positive control, and two variable inputs C3 and Tm1 upon expressing csChrimon, and recorded calcium responses in Tm9. These experiments were done in a *norpA* mutant background, where red light alone did not show any visually evoked responses (data not shown). While activation of L3 elicited the strongest responses in Tm9, activation of all three neurons caused significant responses in Tm9, supporting the notion that these are all functional presynaptic inputs (Fig. 2e). In addition to the stochasticity in the presence of an input, the rank of each input could also vary (Extended Data Fig. 2b). While L3 was the most strongly connected input in most columns, either Mi4 or CT1 were the second most common input. Afterwards, the order became random. For example, the fourth most common input Tm16 could range from being the second to the tenth most common input to a given Tm9 neuron (Extended Data Fig. 2b). Together, these data suggest that there is remarkable variability in the presence, strength and rank order of synaptic connectivity onto Tm9.

### Heterogeneity in presynaptic inputs is not a general feature of all visual system neurons

To understand how much of this variability in connectivity is specific to Tm9 with its unique variable physiological properties, we compared connectivity of Tm9 to that of two other medulla interneurons, Tm1 and Tm2. Like Tm9, Tm1 and Tm2 are essential neurons of the OFF pathway, with similar projection patterns to Tm9^3,31,33,52^. Unlike Tm9, their receptive field widths and temporal kinetics are similar across different columns and across flies ^29^. We identified pairs of Tm9, Tm1 and Tm2 in 166 columns of the right optic lobe (Fig. 3a). As for Tm9, we also mapped the presynaptic input architecture for Tm1 and Tm2, for all neurons with equal or more than four synaptic inputs, together accounting for 80.0 ± 6.0 % of its synaptic partners for Tm1 and 76.3 ± 5.9 % for Tm2 (72.4 ± 8.1 % for Tm9 within this dataset). We identified and annotated the presynaptic inputs of Tm1 to 80.0 ± 6.0 % and Tm2 to 76.3 ± 5.9 % completion.

**Fig. 3:**
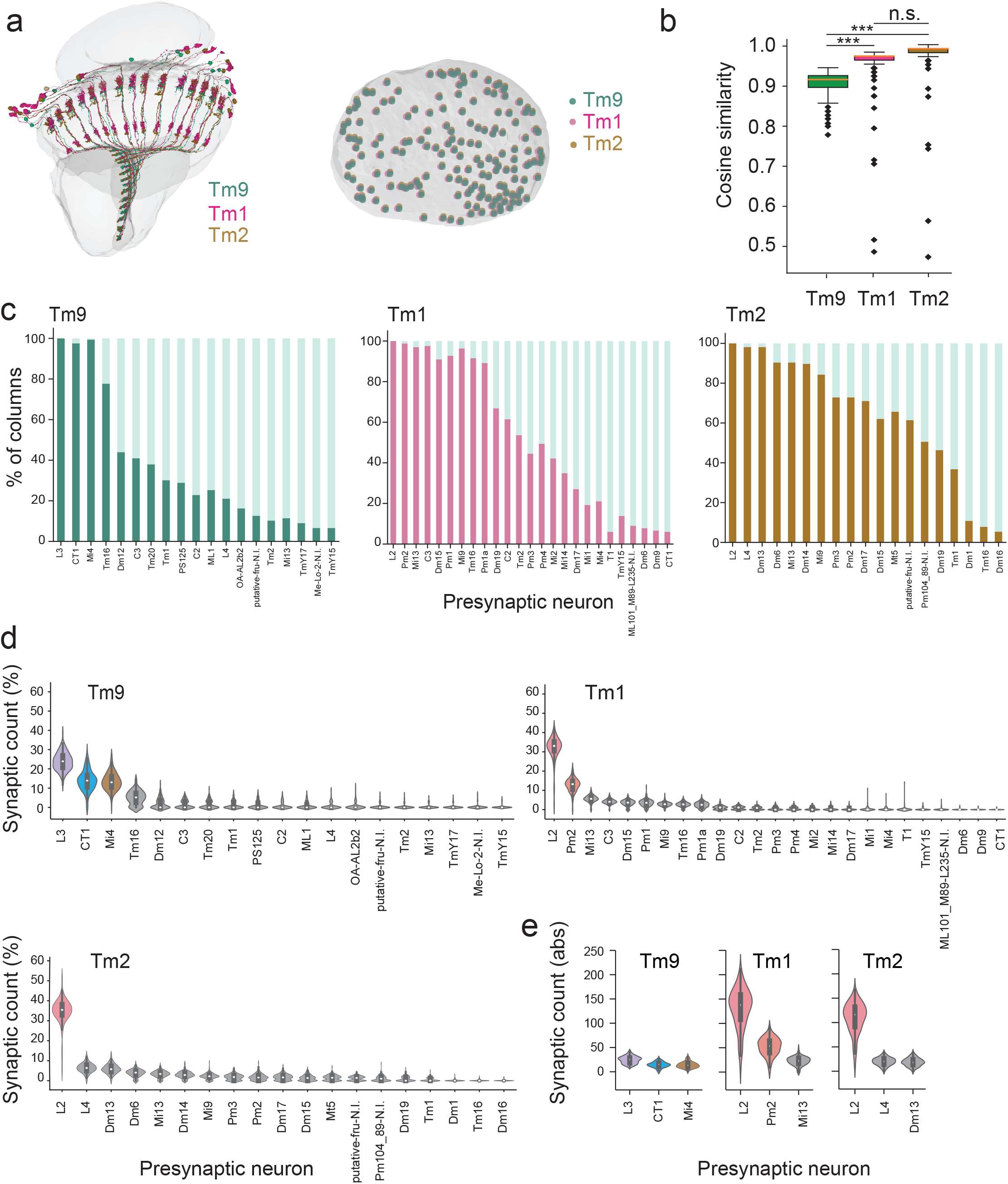
Tm9 has more heterogeneous input partners than other Tm neurons. **a)** Left: Some example Tm9, Tm1 and Tm2 neurons from the same columns. Right: Top view of the medulla, each green/magenta/brown dot depicts one Tm9/Tm1/Tm2 analyzed. **b)** Cosine similarity within Tm9, Tm1 or Tm2 datasets. Statistical testing was done the Kruskal-Wallis after Dunn-Bonferroni correction for multiple comparison *** p<0.001, n.s. = non-significant (p>0.05). **c)** Bar graphs illustrating the % of columns in which a presynaptic input is present for Tm9, Tm1 and Tm2. **d)** Violin plots of the relative synaptic counts of Tm inputs, showing the median (white dot), the interquartile range (thick gray bar), 1.5x interquartile range (thin gray line) and outliers. Neurons with more than 10 % contribution of all synapse counts are color-coded. **e)** Violin plots of the absolute synaptic counts of the first three major inputs to Tm neurons.

Among these three neurons, the connectivity (cosine) similarity was lower within the Tm9 dataset, as within the Tm1 and Tm2 similarity, arguing that presynaptic circuitry of Tm9 is more heterogeneous (Fig. 3b). Although all three Tm neurons received input from a similar number of cell types (Extended Data Fig. 3a), more cell types were prominently present as presynaptic input to Tm1 and Tm2 as compared to Tm9. For example, 12 and 14 cell types gave presynaptic input to more than half the Tm1s and Tm2s, respectively, but only four cell types gave presynaptic input to more than half the Tm9s (Fig. 3c, Extended Data Fig. 3a). In addition to differences in the presence or absence of inputs, there were also differences in synapse counts. Tm1 and Tm2 receive major input from one neuron, L2, whereas, all three non-variable inputs to Tm9 (L3, CT1, Mi4) contributed fairly evenly (Fig. 3d). This difference became even more obvious when looking at absolute synapse counts: whereas L2 made more than 100 synapses with Tm1 or Tm2, L3 only made 23.9 ± 4.5 with Tm9 (Fig. 3e). So far, we compared synaptic connectivity of different presynaptic cell types. To investigate if the variability of synapse counts made by each cell type differs between Tm neurons, we analyzed the coefficient of variation for each presynaptic cell type. This showed that there were no striking differences between Tm9, Tm1 and Tm2 (Extended Data Fig. 3b). Taken together, the variability of synapse counts within each presynaptic column is similar between Tm9 and other Tm neurons. However, Tm9 neurons are more heterogeneous than Tm1 and Tm2 regarding the pure presence of their inputs. Furthermore, whereas Tm1 and Tm2 have one predominant input, the major inputs to Tm9 are more distributed.

### Circuit motifs that can contribute to variable spatial and temporal properties of Tm9

We next asked whether there is some structure within the connections between Tm9 and its presynaptic inputs, and if there are distinct motifs that define variable Tm9 properties. We explored correlations between all Tm9 input counts across the optic lobes, only the two GABAergic neurons Mi4 and CT1 were significantly (anti-) correlated, contributing antagonistically to Tm9 connectivity in different columns (Fig. 4a). An alternative approach, K-means clustering for 6 clusters (inspired by Tm9 physiology, Fig. 1c), resulted in 4 clusters with higher CT1 counts compared to Mi4 counts and 2 clusters vice versa (Extended Data Fig. 4a), suggesting that one of the main sources of variability is coming from the antagonism of CT1 and Mi4. To investigate this further, we re-clustered for two clusters and performed dimensionality reduction using principal component analysis (PCA), analyzing all input counts to Tm9 (Fig. 4b,c). Mi4 and CT1 contributed antagonistically to the two principal components (PC) that explained the most variance (Fig. 4b, Extended Data Fig. 4b,c). Displaying the CT1-dominated and the Mi4-dominated subtypes in PC space revealed a continuous variability since Tm9s from both clusters spanned the PC space continuously rather than forming distinct clusters (Fig. 4c). In the optic lobe, Tm9 neurons belonging to the two clusters were distributed across the medulla (Fig. 4d). Together, these data suggest that two Tm9 subtypes can be defined by the dominated connectivity from one of two major GABAergic inputs, CT1 or Mi4.

**Fig. 4:**
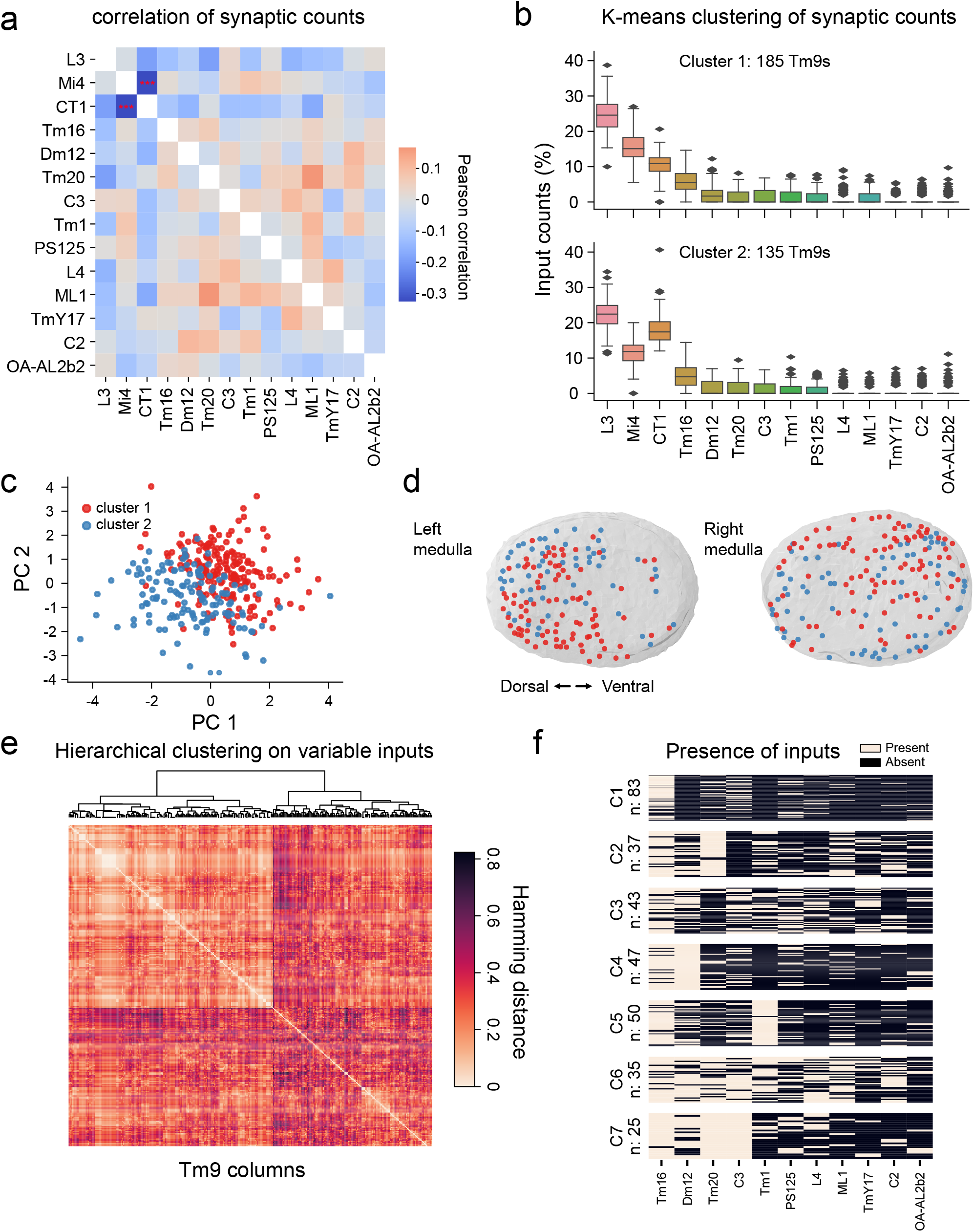
Circuit motifs in Tm9 presynaptic connectivity. **a)** Heatmap of the Pearson correlation between Tm9 input neuron counts. *** p<0.001 after Bonferonni correction for multiple comparisons. **b)** Input counts of Tm9 neurons upon K-means clustering of input connectivity with a cluster number of two. c) Tm9 input connectivity projected onto PC1 and PC2 with labels representing clusters from K-means (cluster 1, red; cluster 2, blue). d) The location of the analyzed Tm9 neurons in the medulla, colored to represent the K-means clusters. e) Heatmap showing the Hamming distances between Tm9 columns ordered based on the dendrogram upon hierarchical clustering considering only the variable inputs (excluding L3, Mi4, CT1). f) Input motifs from the clustering approach in (e). Each row of the binary image represents a Tm9 neuron (n=320), the binary color code indicates the presence or absence of input neurons within each cluster. 7 clusters were chosen based on the silhouette coefficient after clustering with different numbers of neurons (n, given to the left of the plot).

Because neurons with low synaptic input counts can have a profound effect on neuronal function ^53,54^, we next explicitly explored the non-common inputs to Tm9. We binarized the data (a presynaptic cell type being present or absent) and performed Hamming-distance based clustering of the analyzed columns. This analysis revealed several groups, suggesting that they have distinct presynaptic input characteristics (Fig. 4e). We then explored several clusters for the presence or absence of input neurons in order to understand the circuit motifs that contributed to them (Fig. 4f). While Tm16 was present in most clusters, other neurons - especially Dm12, Tm20 and C3 - dominated a cluster by either being present or absent as presynaptic inputs to all Tm9s of that cluster (Fig. 4f). Thus, specific non-common inputs to Tm9 further contribute to forming variable circuit motifs.

### The heterogeneous Tm9 inputs show no spatial structure

What type of neurons are these cell types, and how are these circuit motifs distributed in space? The latter question is especially interesting because previous work had suggested that Tm9s of the dorsal and ventral medulla are developmentally genetically separable ^55,56^. C3 is a GABAergic feedback neuron that, together with its sibling C2 ^5,57^, could regulate temporal properties of their postsynaptic partners (Fig. 5a). When analyzing the spatial distribution of these cells across the dataset, there was no apparent structure. This was true both in the 320 columns analyzed across the two optic lobes, as well as in a dense patch, in which we analyzed each column (n = 57) of the visual system in order to look at spatial structure between neighboring columns (Fig. 5b). Another cell type that contributed to presynaptic circuit motifs, Dm12, is an amacrine-like wide field neuron that spans several columns ^44^. Such neurons could pool information from several columns before passing it on to Tm9. This is interesting with respect to the variability in the Tm9 spatial receptive fields. Both Dm12, and the other wide field neuron in the dataset, Tm16, are thus candidates to widen the size of the Tm9 receptive field (Fig. 5c). Interestingly, Dm12 and Tm16 did not only vary in their presence or synaptic counts to Tm9, but also in how many cells of these types are connected to one Tm9, a number that can vary from 0 to 4 (Fig. 5c,d). In addition to the columnar feedforward and feedback neurons, and the multi columnar neurons identified here, other neurons could still further contribute to functional variability of Tm9. We for example identified all OA-AL2b2 neurons of the fly brain ^46^ as input to Tm9 (Fig. 5e). These are four giant neurons, two per hemisphere, which are either octopaminergic or cholinergic ^46,58^ and extend thin projections across the entire medulla. Several Tm9s are connected to each of these neurons. Such global neurons could for example modulate physiological properties of its postsynaptic partners based on the physiological state of the animal ^29,59^. Interestingly, 13% of all neuronal projections from the central brain target the optic lobe ^41^, suggesting that such feedback is widespread and prominent. Overall, we found no apparent spatial organization for any of the heterogeneous Tm9 inputs, arguing that their distribution across the medulla is stochastic.

**Fig. 5:**
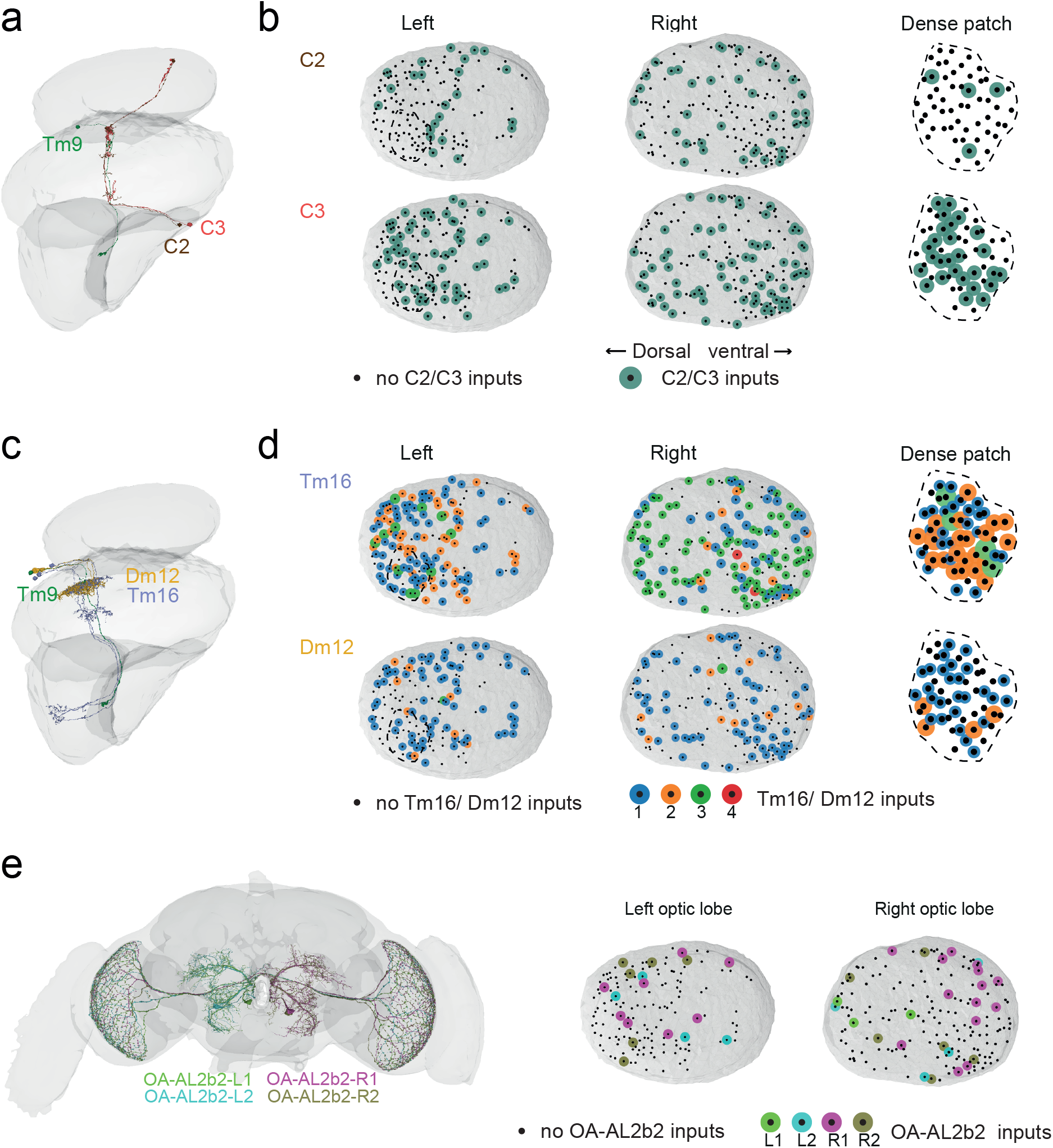
Different types of cells that connect to Tm9. **a)** Illustrations of one Tm9 (green) with its input partners C2 (brown) and C3 (red). **b)** Top view of the medulla with each black dot depicting one Tm9 neuron analyzed, green dots show the presence of C2 (top) or C3 (bottom) as an input partner to these Tm9s for the left (left) and right (middle) optic lobes. The same is shown for the dense patch, a region in the left optic lobe (dotted lines) in which Tm9s were analyzed in each column. **c)** Illustrations of one Tm9 (green) getting input from two Dm12s (yellow) and three Tm16s (blue). **d)** Same as in b) for Tm16 and Dm12, only that the color of the dots represents the number of cells of one type connected to Tm9 in that column. **e)** Image of all four OA-AL2b2 cells (left) as well as their individual connectivity to Tm9.

### Heterogeneity of Tm9 connectivity exists across flies

The analysis presented here, and connectomics analysis in general, have one major limitation: data are generated from an individual specimen. The FAFB dataset is the first EM volume that allows a comprehensive analysis of visual circuitry across the fly eye. To ask whether the heterogeneous synaptic Tm9 inputs identified in the FAFB dataset are heterogeneous Tm9 inputs in general, we used expansion microscopy (ExM) to study wiring variation across individual animals ^60,61^. To achieve specificity between two cell types of interest, we made use of the genetic toolkit of the fly and cell type-specifically labeled presynaptic sites using the active zone marker Brp[short]::mCherry and postsynaptic dendrites using membrane-tethered GFP (Fig. 6a,b). After imaging the expanded optic lobes, we automatically counted the presynaptic sites directly opposing a Tm9 dendrite across many columns in several flies per genotype based on previously described distance measurement between Brp-marked active zones and the postsynapse (Fig. 6a,c)^62^. This analysis broadly reflected the results obtained from the FAFB dataset for three cell types tested, Tm16, Dm12, and C3. For example, Tm16 was connected to Tm9 in most columns and only a few columns did not show any Brp-positive puncta for Tm16 - Tm9 pairs (Fig. 6c,d). The distribution of synapse counts were also similar between the one connectome and genetically labeled optic lobes (Fig. 6d). Light microscopy data only slightly underrepresented absolute synapses numbers as compared to the EM dataset. This is likely attributable to the fact that active zones that are very close together still cannot be resolved by light-microscopy based measurements. Taken together, this data supports the idea that specific hypotheses based on EM data can be tested using the approach taken here ^60,63^. Furthermore, heterogeneous presynaptic partners of Tm9 as suggested by EM could be verified to provide heterogeneous input to Tm9 in general. This correlates with the observation that heterogeneous physiological properties of Tm9 have been reported within as well as across flies ^64^ (Fig. 1c).

**Fig. 6:**
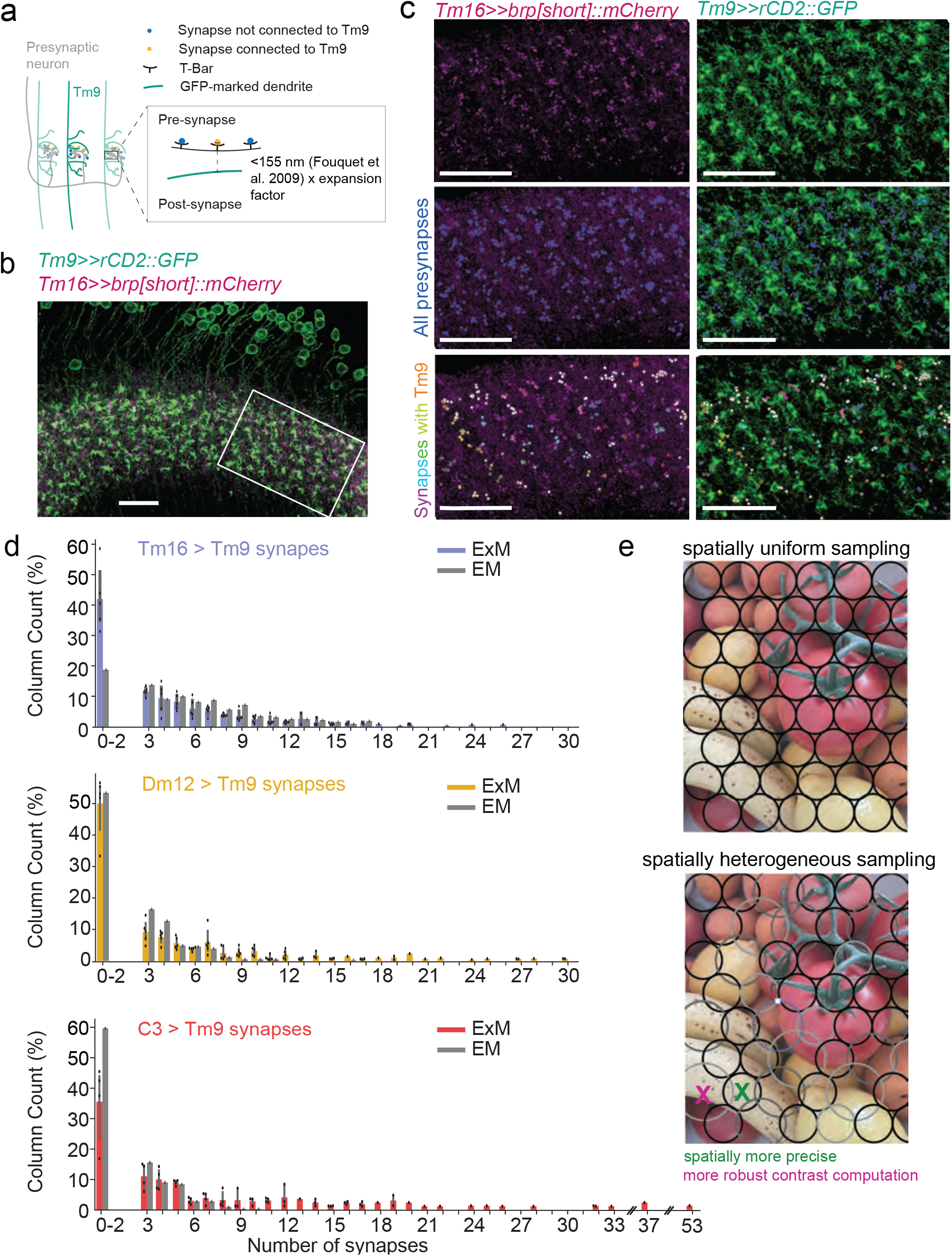
Expansion microscopy confirms heterogeneity of presynaptic inputs across flies. **a)** Schematic of the experiment. **b)** Confocal image of a medulla in which Tm9 dendrites are marked with rCD2::GFP (green) and Tm16 presynapses with Brp[short]::mCherry (magenta). **c)** Confocal images showing the boxed area in **(b)**. All presynapses are identified (blue, middle), and the ones in close proximity to Tm9 dendrites are sorted into columns (different colors, bottom). All scale bars are 20 μm **d)** Barplots showing the percent columns in which a specific presynaptic neuron (Tm16, Dm12, and C3) makes a certain number of connections with Tm9. Bar graphs of ExM data show 95 % confidence interval and all individual data points. **e)** A visual scene as sampled by a columnar visual neuron with uniform spatial receptive fields (top) or by a columnar visual neuron with spatial receptive fields of various sizes (bottom). Small receptive fieldsare spatially more precise (green), whereas bigger receptive fields can contribute to more robust contrast computation (green).

## Discussion

Visual systems have been considered homogeneous structures. Here, we describe an exception to this rule. We analyze a specific circuit in the *Drosophila* visual system, centered around the visual interneuron Tm9, and describe that this circuit shows heterogeneity in synaptic connectivity. In comparison to Tm neurons, which also show some variability in their presynaptic wiring, the variability is more extensive for Tm9. Certain circuit motifs exist, which can set the basis for physiological variability described for this cell type, but without any apparent spatial structure, arguing that the distribution of the presynaptic input structure across the visual system is stochastic.

Based on most criteria that have been used to identify cell types, Tm9 still counts as a prototypic cell type. It is anatomically distinct, and easily distinguishable from all other cell types in the visual system ^5^. It is a columnar neuron whose dendrites tile the visual system. And it is easily distinguishable by distinct genetic markers, allowing very specific genetic access ^31^. Cell type assignments have long relied mostly on anatomical criteria, supported by genetic markers, and physiological criteria have rarely served alone to define a cell type ^10^. However, recent population imaging approaches have classified cell types in the vertebrate retina based on functional properties ^65–67^. These studies found more types based on function as compared to other criteria ^66^. This, together with our work here argues that functional analysis alone can serve to define many types, but if cell types with heterogeneous properties such as Tm9 are present, purely functional analyses might overestimate the true number of cell types. To comprehensively distinguish all cell types present, one will still need to consider a diverse set of criteria, and potentially apply them flexibly.

Visual systems are considered to be built by genetically deterministic mechanisms. For example, genetically encoded sources of variation, especially gradients of signaling molecules, play a prominent role in developmental biology. Recent transcriptomics studies in fact revealed two subtypes of - dorsal and ventral - Tm9, distinguishable by the developmental expression of Wnt10 and Wnt4 ^55,56^. However, our data showed no evidence for differences in connectivity between the dorsal and ventral halves of the optic lobe. Furthermore, specific molecular mechanisms achieve layer- and column-specific targeting that results in the repetitive structure of the eye ^6^. Our understanding of an increasingly intricate connectivity of visual neural networks challenges this view, and additionally complexity in the presence or number of synaptic partners extends beyonds the limits of the capabilities of genetic determinism. Recent discoveries that an interplay between stochastic presynaptic filopodial dynamics and the relative availability of postsynaptic partners controls synapses formation ^23,24^, provide an attractive mechanism by which non-identical, stochastic patterns of connectivity across the eye could arise. Connectivity pattern could be truly random, or could be shaped by spontaneous or activity-dependent plasticity to achieve an appropriate behavioral output. Additional environmental differences can further contribute to variability across animals ^68^.

What are the potential advantages of such heterogeneous properties? Tm9 is an interesting neuron, because it plays an essential role in the establishment of direction-selective signals, a computation that requires the comparison of local luminance changes over space and time ^29–33,51^. At the same time, its major presynaptic partner L3 has been shown to be luminance-sensitive and to mediate rapid luminance gain control in a quickly changing environment ^69–71^. Here, it could be advantageous to integrate luminance over not-so-local regions of visual space, and also over time, to correctly estimate the background luminance of the visual scene. Thus, Tm9 neurons with narrow spatial receptive fields or fast response properties might have an advantage for detecting local motion cues, whereas Tm9 neurons with bigger spatial receptive fields and slower, more integrating properties might be better at adjusting luminance gain, which in turn leads to a more accurate computation of contrast (Fig 6e). The distribution of neurons with diverse properties across the optic lobes argues for a division of labor. Given that most, if not all, visual cues relevant for behavior stimulate larger regions of the eye, stable behavior might tolerate such a distribution of tasks over different columns. In fact, distributing tasks within one cell types (Fig. 6e) might actually be advantageous, for example by being less costly. In the future, the availability of a first full connectome of the fly visual systems will allow investigating how widespread this variability is across visual cell types. Ultimately, it will be interesting to understand how wiring variability within the visual systems contributes to robust circuit function, or to individuality of behaviors.

## Author contributions

MS, JCo, and SMO conceptualized the work, SMO, AB, JCh, LL, MI, BG, RT and MS performed analysis in FlyWire, SMO, JCo and BG analyzed connectome data, JCo and GRT collected and analyzed imaging data, MS and CS supervised and acquired funding, MS and JCo wrote the initial draft of the manuscript, SMO, CS, AB and BG edited the manuscript.

## Acknowledgments

We thank Michael Reiser and Henrique Ludwig for training on the FAFB dataset, and many insightful discussions, Annalena Oswald for help with proofreading, Jonas Peper for technical help with establishing expansion microscopy, and Philipp Schlegel helpful input on FlyWire analysis. We are grateful to all FlyWirers who contributed to the proofreading and annotation of optic lobe neurons. This project has received funding from the European Research Council (ERC) under the European Union’s Horizon 2020 research and innovation program (grant agreement No 716512) to MS, and from the German Research Foundation (DFG) through the research unit FOR 5289 (project P5) to MS, as well as DFG grant SCHN1479/1-1 to CS. RT acknowledges a PhD stipend by the FTN Mainz.

## Additional information

Supplementary information is available for this paper.

Correspondence and requests for materials should be addressed to Marion Silies.

## Competing interests

The authors declare no competing interests.

## Materials and Methods

### EM data analysis

We analyzed connectivity of Tm neurons (Tm9, Tm1, Tm2) using the EM dataset of a full adult female brain (FAFB) ^39^. We initially identified Tm9 neurons in the dataset by making use of the already annotated CT1 neuron, which was known to be postsynaptic to Tm9 in each medulla neuron column ^32^, we here found that it is also a common presynaptic partner of Tm9. We identified further Tm9 neurons by exploring regions in the medulla or lobula plate neighboring Tm9 at similar levels as Tm9 axon terminals or Tm9 dendrites. To identify Tm1 and Tm2, which project to the same lobula plate layer as Tm9, we explored segments in close vicinity to previously identified Tm9s. We then proofread all Tm neuron backbones. We also performed more detailed, ‘twig proofreading’ for 16 Tm9 and 9 Tm1 neurons, which did not lead to significant changes in synapse number.

Using the Buhman algorithm for automated synapse annotation ^43^ we acquired neuron segments corresponding to the presynaptic partners of Tm neurons using a cleft score >= 50 (as in ^42^). We filtered out all redundant synapses if they were less than 100 nm apart. To interact with the human-AI reconstruction platform FlyWire (https://flywire.ai/) programmatically, we employed the python package fafbseg (https://fafbseg-py.readthedocs.io/en/latest/source/api.html).

After selecting Tm9 neurons in many medulla columns distributed across both optic lobes, we excluded columns in which the lamina input L3, the major input of Tm9 ^30,31^, had a darkened cytoplasm in the EM micrographs. This was most prominent in the part of the left optic lobe in which the lamina was detached, but also along the edges of the optic lobes. In these areas, cells with a disrupted membrane were darker, most likely due to higher exposure to the fixative, and not due to biological damage. We conclude this, because we were able to manually identify ‘normal looking’ synapses within these neurons, while the Buhman algorithm failed to recognize the synapses.

We aimed to identify ∼80 % of all inputs to a given Tm neuron. Based on this, we traced and annotated presynaptic segments making >= 3 synapses (Tm9) or >= 4 synapses (Tm1, Tm2). Synapse counts below these thresholds were set to 0.

#### Annotation of cell types

Many cell types in the visual system have been described. Here, cells were first assigned to a certain type based on anatomical similarities with cell types described previously ^5,19,36,37,44–46^. If we could not easily assign a cell to one type, plotting several cells of one potential type usually helped to verify or falsify a guess based on overlap in the branching patterns and similarity in the overall structure of neurons. This was especially true for cells at the edges of the optic lobes, where dendritic organization wasn’t as clear as in the center of the visual system. All cell type annotations not done by us, but by other members of the FlyWire community, were verified using the same method. For cells that were hard to distinguish we further queried connectivity of cells under question using Codex (flywire.codex.ai). Cells with elaborate projections into the central brain were classified based on a comparison with the hemibrain dataset ^38^ using NBLAST ^47^.

We only included neurons in the analysis which had connections to Tm9 in more than 5 % of the columns.

### Data analysis of FAFB data sets

Custom-written code in Python version 3.9 was used to further process data, including the Python packages pandas (https://pandas.pydata.org/), numpy (https://numpy.org/), scipy (https://scipy.org/), and matplotlib (https://matlplotlib.org/) for analysis, statistics, and data visualization. The flywire.ai web interface or Navis python packages were used for anatomical visualization. For visualization, data were sorted by mean count per presynaptic cell type.

To define spatial coordinates for each cell, we obtained the center of mass of all postsynaptic sites of Tm9 in its dendritic region. The closest presynaptic site to this center was taken as the spatial coordinate for that cell. Tm9 neurons were assigned to the dorsal (D) or ventral (V) medulla after projecting all 3D-Tm9 coordinates to a 2D plane using Singular Value Decomposition, and drawing a midline at the centroid of this shape.

A correlation matrix (Fig. 4a) was calculated using the pairwise Pearson correlations of Tm9 synaptic input neuron counts. To determine significance in multiple comparisons, we corrected the α using the Bonferroni method. K-means clustering was performed using the scikit-learn Python package with either 2 (Fig. 4b) or 6 clusters (Extended Data Fig. 4b). Principal components analysis was performed by first standardizing the Tm9 relative input counts and computing the eigenvectors of the covariance matrix of the data. Then the data were projected onto two principal components using the two eigenvectors with highest eigenvalues. For hierarchical clustering of Hamming distances, we first binarized the Tm9 input counts (presence or absence of neurons) and calculated the pairwise Hamming distances of Tm9 columns using the scikit-learn Python package. Hamming distance captures the number of different bits between binary data, providing an interpretable measure of similarity between Tm9 columns. We then used agglomerative clustering (scikit-learn) to perform hierarchical clustering and determined the optimal number of clusters as 7 by maximizing the silhouette coefficient between 4-15 clusters (a range of expected number of input motifs based on physiological data).

### Fly husbandry and genetics

All *Drosophila melanogaster* were raised at 25 °C and 65 % humidity on molasses-based food while being subjected to a 12:12 hour light-dark cycle.

For expansion microscopy, 3-9 day old female flies were used. The driver line for the postsynaptic cells was *Tm9*^*24C08*^*-LexA* (BL 62012), recombined to *UAS-brp*^*short*^*::mCherry*. The line w+; *Tm9*^*24C08*^*-LexAp65*^*attP40*^. *UAS-brp*^*short*^*::mCherry/* CyO; *LexAop-rCD2::GFP/TM6B* was crossed to different drivers for the presynaptic inputs: *Tm16*^*82C05*^*-Gal4-AD*^*attP40*^*;Tm16*^*52D11*^*-Gal4-DBD*^*attP2*^ (SS00335 Aljoscha Nern), *Dm12-split-Gal4* (SS00359, Aljoscha Nern, Janelia Research campus) and *C3*^*R26H02*^*-Gal4-AD; C3*^*R29G11*^*-Gal4-DBD* (Tuthill et al. 2013).

Calcium imaging experiments used female flies, 1-7 days post-eclosion of the genotype *w/w+; Tm9*^*24C08*^*-LexA,lexAop-GCaMP6f/+;+/+*. For optogenetics experiments, fly food was supplemented with all-trans retinal (ATR) to reach a concentration of 1 mM, as in (Hornstein, Pulver, and Griffith, 2009 REF), *Tm9*^*24C08*^*-LexA,lexAop-GCaMP6f* were put into blind, *norpA*^*36*^ mutant flies, and *norpA* mutant males were imaged. *L3*^*0595*^*-Gal4* ^48^, *Tm1*^*27b*^*-Gal4* ^52^ and the C3-split-Gal4 line *R35A03-AD;R29G11-DBD* were used to excite L3, Tm1, or C3 neurons.

### Expansion Microscopy

Flies were anesthetized on ice and dissected in 1x phosphate-buffered saline (PBS, Gibco Thermo Scientific). For better positioning during mounting, the optic lobes were separated from the central brain and collected on ice in Terasaki plates. After keeping them on ice for no longer than 20 minutes they were fixated for 50 minutes in 2 % paraformaldehyde (PFA, Polysciences, diluted in PBS) while shaking at room temperature (RT). The optic lobes were washed three times in PBT (PBS with 0.3 % Triton x-100, Roth) for 10 minutes, then blocked in 10 % normal goat serum (NGS, Thermo Scientific, in 0.3 % PBT) for 50 minutes, both at RT while shaking. Afterwards, the samples were incubated in primary antibody (chicken anti-GFP, Abcam #13970, 1:2000, rabbit anti-DsRed, Takara Bio Clontech #632496, 1:200) for 24 – 48 hours in the dark while shaking at 4 °C, then washed three times in PBT and incubated in secondary antibody mix (goat anti-chicken Alexa Fluor 488, Jackson Immunoresearch #103-545-155, 1:200, goat anti-rabbit ATTO 647N, Sigma #40839, 1:100) for 24 - 48 hours in the dark while shaking at 4 °C. Optic lobes were then washed 2x in PBT and 2x in PBS before either mounting them in Vectashield (Biozol) for unexpanded brain imaging or continuing with the expansion microscopy protocol.

The expansion microscopy protocol was modified after ^61^. Optic lobes were anchored in Acryloyl-X, SE (Sigma, 1:100 in PBS) over night at 4 °C in the dark while shaking and then washed 3x with PBS for 5 minutes in the dark on ice. For polymerization, the monomer solution (7.4 % sodium acrylate, 2.5 % acrylamide, 0.15 % N,N’-methylenebisacrylamide (all Sigma), 2 M Sodium Chloride (Roth), 1x PBS) was prepared on ice. Before the brains were incubated in the monomer solution, 0.1 % ammonium persulfate (Honeywell), 0.1 % tetramethylethylenediamine (TEMED, Roth) and 0.005 % 4-hydroxy-2,2,6,6-tetramethylpiperidin-1-oxyl (4-hydroxy-TEMPO, Sigma) were added to activate the polymerization reaction. Brains were placed three times in fresh monomer solution, incubating for five and 20 minutes on ice in Terasaki plates. For the last step the samples were transferred onto a gelation chamber, a glass slide (Menzel-Gläser) with two cover glasses (Marienfeld, 18x18 mm, No.1) as spacers. The lobes were positioned preferably with the cut side down in monomer solution and covered with a cover slip. Afterwards, the space between spacers was filled with monomer solution to prevent drying of the gel. The filled chambers then were incubated for 1.5 hours at 37 °C. After a gel was formed with embedded tissue the excess gel was cut off. The remaining gels with the specimens were digested each in 1 ml digestion buffer (50 mM Trizma pH 8.0 (Roth), 0.5 % Triton X-100, 0.8M Guanidine HCl (Sigma), 1 mM EDTA pH 8.0 (Roth) and 1:100 Proteinase K (New England Biolabs 20 mg/ml)) in a 2 ml Eppendorf tube for at least 24 hours at RT in the dark. The gels were then carefully washed and expanded with deionized water, three times for at least 15 minutes. They were carefully poured onto a cover glass (Marienfeld, 24 × 60 mm, No.1) with a 3D printed frame (heights 0.3 mm) glued to it. The excess water was removed, small amounts of 2 % low melting point Agarose (Sigma) was pipetted to the edges of the gel to reduce movement and Vectashield was used as a mounting medium before covering the samples with a glass slide. The optic lobes were imaged on a Leica Stellaris 8 STED microscope equipped with a 93x objective.

### Analysis of Expanded Samples

To reduce background noise the VVDViewer for large volume segmentation ^63^ was used. Both the *Tm9>>rCD2::GFP* channel and the Brp signal channels were treated separately. Intensity thresholds were individually set for each brain so that the background was reduced, and the dendritic arborizations of *Tm9>rCD2::GFP* signal were well visible. Subsequent analysis was done with custom-written code using the python package napari (Python 3.9). Local maximum detection identified Brp puncta within the Brp^short^::mCherry signal. To eliminate noise, a distance threshold of 200 nm between two Brp puncta was used based on findings from Gao et al. 2019 who reported that this was the distance between separable Brp signals. To quantify the number of synapses per column, columns were manually identified using an napari-ROI selection tool (Windhager 2023) around single Tm9 cells. Only for *Tm9>>rCD2::GFP, C3>>brp*^*short*^*::mCherry* the Brp channel was used to draw ROIs around clearly visible columns formed by the *C3>>brp*^*short*^*::mCherry* signal. After defining single columns the GFP mask was binarized. This mask was used to calculate the minimum distance to each Brp puncta. By using a distance threshold of 300 nm (value after ^62^, multiplied with the expansion factor) the Brp puncta closest to the *Tm9>>GFP* mask were detected and counted as synapses.

### In vivo calcium imaging

Flies were immobilized on ice for dissection, and mounted into a sheet of stainless steel foil such that the eyes and rest of the body remained below the foil. The thorax and left head were glued to the foil by UV-cured glue (Bondic). Cuticle, fat bodies, and trachea from the right optic lobe were removed with a razor blade and forceps. This was done under ice-cold, low-calcium saline, which was exchanged with calcium saline at room temperature after dissection. Saline for calcium imaging consisted of 103 mM NaCl, 3 mM KCl, 5 mM TES, 1 mM NaH2PO4, 4 mM MgCl2, 1.5 mM CaCl2, 10 mM trehalose, 10 mM glucose, 7 mM sucrose, and 26 mM NaHCO3 (no calcium, no sugars for low-calcium saline). The perfusion solution was bubbled with carbogen (95 % O_2_, 5 % CO_2_).

Imaging was done using a Bruker Investigator two-photon microscope (Bruker, Madison, WI, USA) coupled to a tunable laser (Spectraphysics Insight DS+). The microscope was equipped with a 25×/1.1 water-immersion objective (Nikon, Minato, Japan). Laser excitation was tuned to 920 nm, and less than 20 mW of excitation was delivered to the specimen, measured at the objective.

Emitted light passed through a SP680 short-pass filter, a 560 lpxr dichroic filter and a 525/70 filter. PMT gain was set to 855 V. The microscope was controlled with the PrairieView (5.4) software. Images of approx. 90 × 256 pixels were recorded at 8–12 Hz, at an optical zoom of 8× to 10×.

Visual stimuli were programmed in C++ using OpenGL and projected onto a rear projection screen (8 × 8 cm, or about 60° × 60° (azimuth × elevation), using a LightCrafter 4500 DLP (Texas Instruments, Dallas, TX, USA) with only blue LED illumination, which was attenuated with a 482/18 bandpass and ND1 filters. Stimulus frames depth was 6-bit pixel, with an update rate of 100 Hz, although the projector frame rate was 300 Hz. The stimulus and data acquisition computers were linked via a NI-DAQ USB-6211 device (National Instruments).

ON-OFF-fullfield flash stimuli covered the whole stimulation screen (60 deg x 60 deg). Stimuli consisted of 2 s OFF and 2 s ON contrast at 100 % Weber contrast, interleaved by a gray background with a duration of 4 s.

Optogenetic activation was done with a 625 nm red LED (ThorLabs). The stimuli consisted of a 25 ms long train of light pulses at 40 Hz. A single pulse of 25 ms was used with a power of 29.40 μWmm^-2^. The pulse started 10 s after the beginning of the recording. Five pulses separated by 30 s from each other were presented per recording.

### Analysis of in vivo calcium imaging data

Data processing was done using custom code written in MATLAB. Imaging time series were registered to compensate for within-plane motion of the specimen, using either cross-correlation alignment or RASL (robust alignment by sparse and low-rank decomposition, by ^72^. To align multiple time series of the same recording plane, a within-time-series registration was followed by an across-time-series registration of the mean of each registered time series that defined the global shift to apply to all frames of the within-time-series registered frames.

Fluorescence time series *F*(*t*) were high-pass filtered with a cutoff period of ≈ 0.1 Hz (or 150 data frames). ROIs were selected manually following the stereotypical shapes of the recorded neuron types. Pixels were averaged within each ROI. The time series was normalized as ΔF/F0=(F−F0)/F0, where the baseline fluorescence *F*0 was chosen as the average fluorescence during all presentations of the background stimulus. To reduce large fluctuations for recordings with baseline signal close to 0, the mean fluorescence of the full time series was added to the denominator `ΔF/F0=(F−F0)/(F0+(Fmean)`). Stimulus and imaging timing were aligned and trial-averaged. Then time series were interpolated to 10 Hz to allow averaging signals acquired at different frame rates.

To visualize the diversity of time courses of the response to contrast changes in Tm9 neurons, we first normalized each trace by subtracting its mean, and dividing by its standard deviation, resulting in the Z-score of the response trace per ROI. The normalized traces have an average of 0 and standard deviation of 1. After removing the influence of the response amplitude, the responses revealed a diversity of time courses. Dimensionality reduction by t-SNE was done to visualize a two-dimensional representation of the response diversity. To visualize this variability, k-Means clustering was performed with the correlation distance between the z-scored responses with a cluster number of 6 which upon visual inspection separated the data best. Cluster assignments were used to color code the z-scored traces to display their different time courses.

## Extended Data

**Extended Data Figures 1-4**.

## Notes

### Competing Interest Statement

The authors have declared no competing interest.

