## Extended Data Figures for "Heterogeneity of synaptic connectivity in the fly visual system"

**Extended Data Figures 1-4.**

### Supplemental Material

**Supplemental Table 1. Contributions to the dataset used in this study.** Shown are the neuron types and numbers annotated by each FlyWirer, and the number of total annotations.

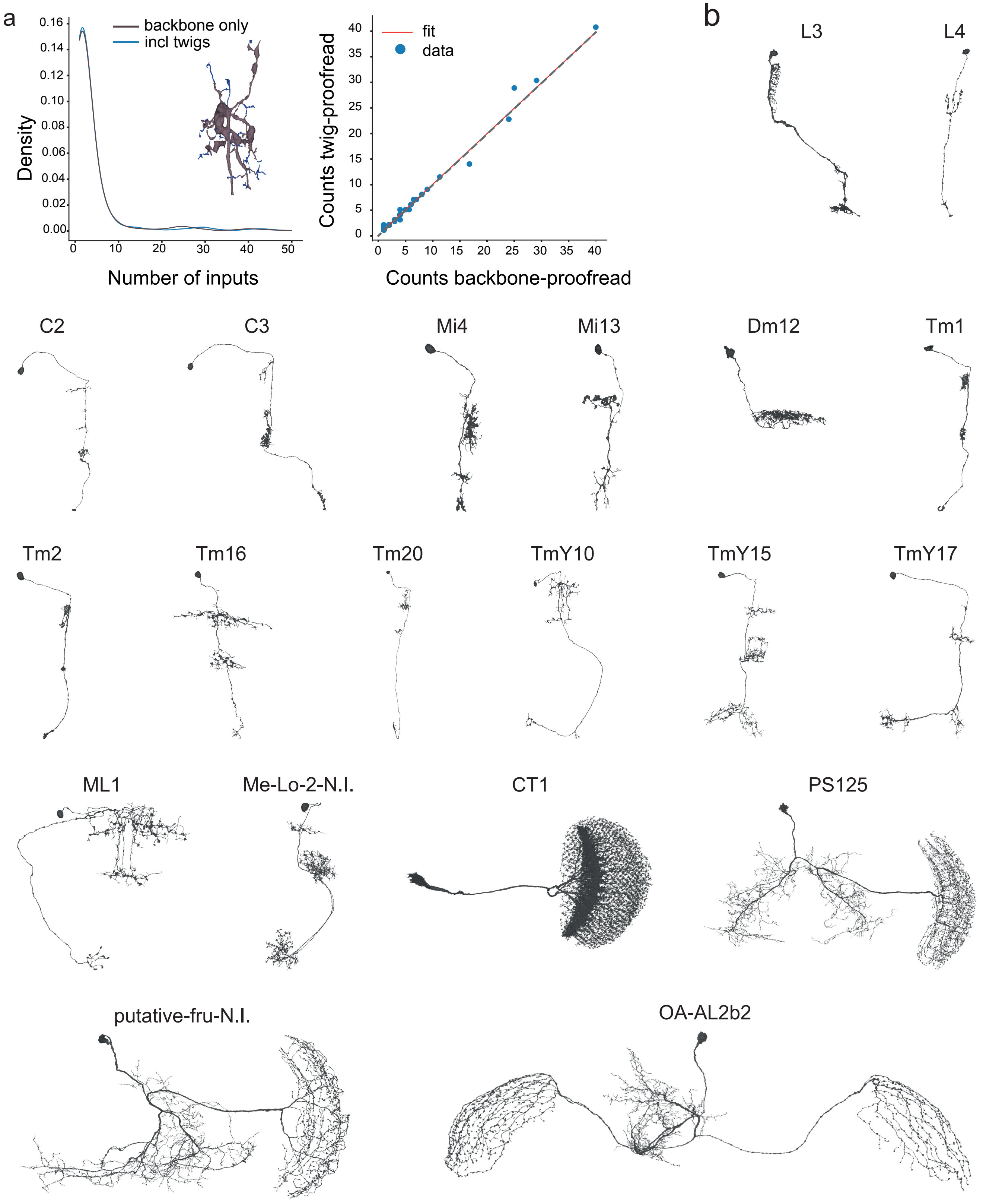


**Extended Data Fig. 1: The presynaptic inputs of Tm9. a)** Left: Density distribution of Tm9 inputs before proofreading. Right: A quantile-quantile (Q-Q) plot depicting the distribution of synaptic connection counts before and after twig-proofreading for Tm9. Each point on the plot represents a pair of quantiles: before twig-proofreading on the horizontal axis, and after twig-proofreading on the vertical axis. The red line represents a linear fit, the gray dotted line is the unity line. **b)** Illustrations of all neurons identified here as presynaptic inputs to Tm9 in >5 % of all columns, micrographs are taken from flywire.ai.

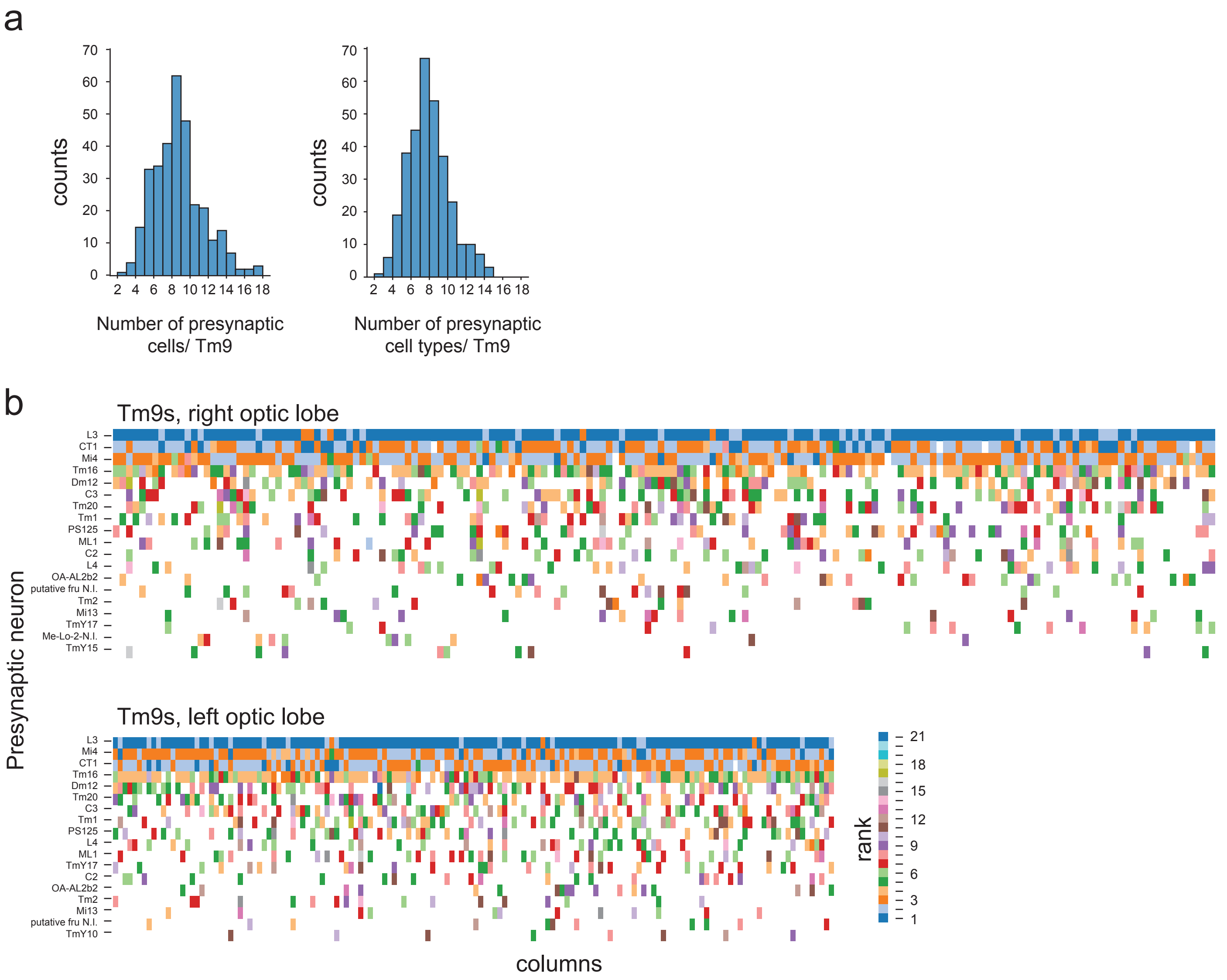


**Extended Data Fig. 2: Analysis of Tm9 presynaptic heterogeneous inputs.**
**a)** Histograms showing how many presynaptic cells (left) and presyanptic cell types (right) individual Tm9 neurons are connected to. **b)** Matrices showing rank of presynaptic inputs to Tm9 neurons of the right (170 columns) and left (150 columns) optic lobes respectively, as long as they are present in >5 % of columns.

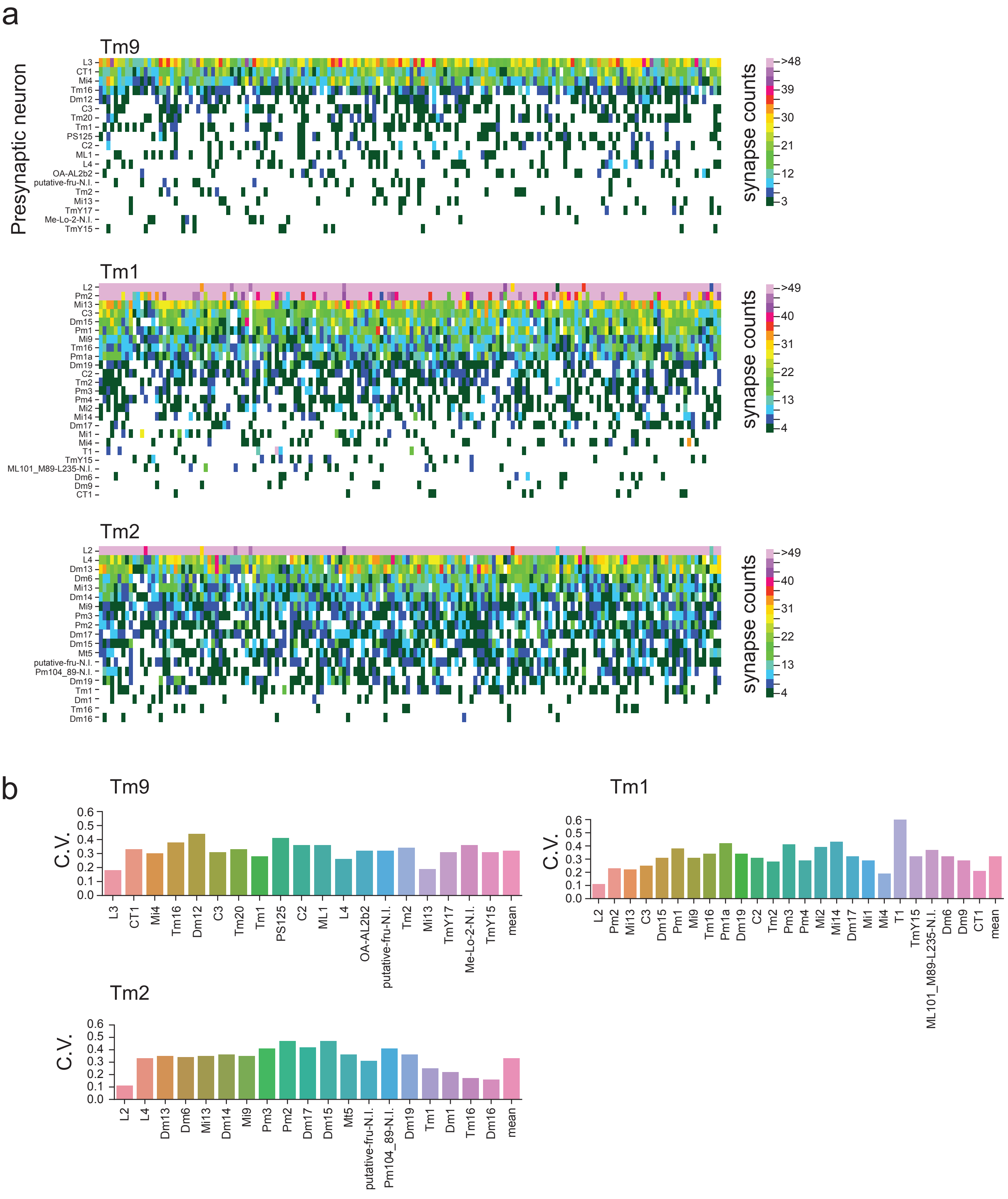


**Extended Data Fig. 3: Comparison of Tm9, Tm1 and Tm2 connectivity. a**) Matrix showing synaptic counts of presynaptic inputs to Tm9, Tm1 and Tm2 neurons of 166 columns of the right optic lobe. **b**) Bar graphs showing the coefficient of variation (c.v.) of relative synapse counts of all the Tm inputs.

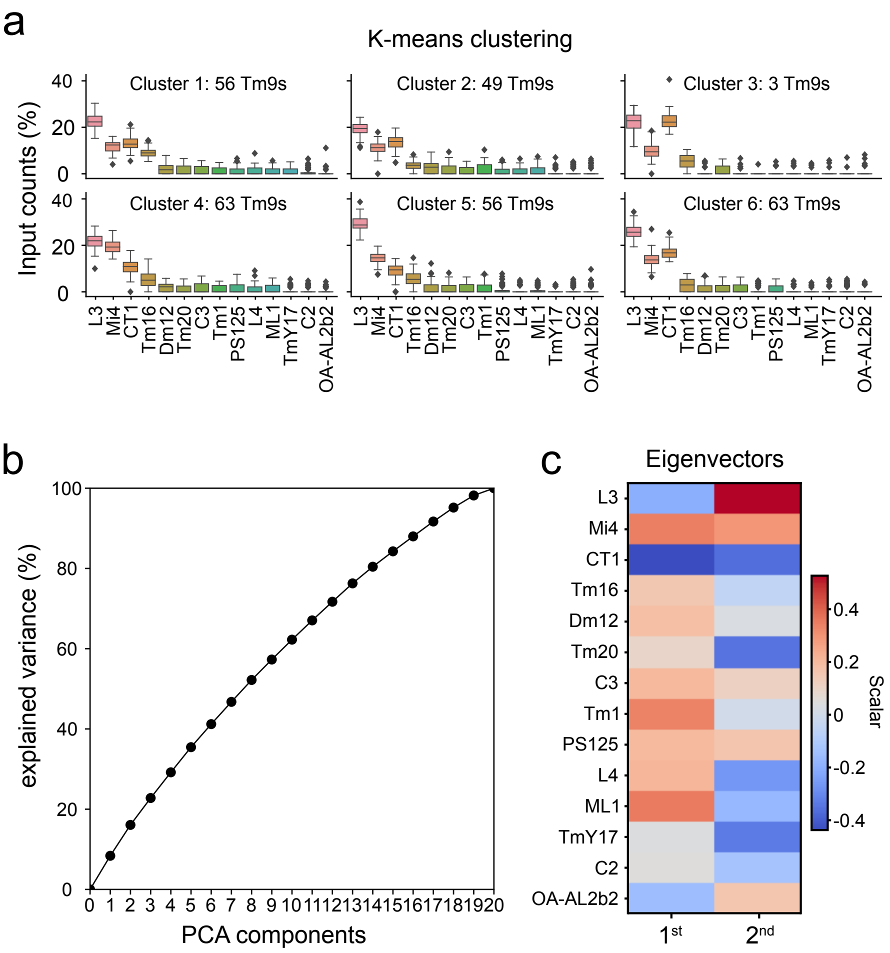


**Extended Data Fig. 4. Analysis of Tm9 circuit motifs. a)** Relative nput counts of Tm9 neurons upon K-means clustering of input connectivity with a cluster number of 6 picked based on physiological clustering. **b)** Explained variance of principal components upon Principal Components Analysis of Tm9 input connectivity. **c)** Contribution of input neurons to the eigenvectors of the covariance matrix of Tm9 input connectivity. The two eigenvectors with the highest eigenvalues which are used to project the dataset in Figure 4c are shown.
